# Modeling the dynamics of EMT reveals genes associated with pan-cancer intermediate states and plasticity

**DOI:** 10.1101/2024.10.03.616309

**Authors:** MeiLu McDermott, Riddhee Mehta, Evanthia T. Roussos Torres, Adam L. MacLean

## Abstract

Epithelial-mesenchymal transition (EMT) is a developmental cell state transition co-opted by cancer to drive metastasis and resistance. Stable EMT intermediate states play a particularly important role in cell state plasticity and confer metastatic potential. To explore the dynamics of EMT and identify marker genes of highly metastatic intermediate cells, we analyzed EMT across multiple tumor types and stimuli via mathematical modeling with single-cell RNA-sequencing (scRNA-seq) data. We identified pan-cancer genes consistently expressed or upregulated in EMT intermediate states, most of which were not previously annotated as markers of EMT. Using Bayesian parameter inference, we fit a simple mathematical model to scRNA-seq data, revealing tumor-specific transition rates. This mathematical model offers a framework to quantify EMT progression. A consensus analysis of differential intermediate expression, regulation, and model-derived dynamics identified marker genes associated with persistence of the intermediate EMT state. *SFN* and *NRG1* emerged as genes with the strongest evidence for their role influencing intermediate EMT dynamics. Through analysis of an independent cell line, we verified the role of *SFN* as a marker intermediate EMT transition. Modeling and inference of genes associated with EMT dynamics offer means to find biomarkers and to identify therapeutic approaches to harness or reverse tumor-promoting cell state transitions driven by EMT.

## 1 INTRODUCTION

Cell state transitions are phenotypic changes in the state of a cell, primarily driven by transcriptional programs. Such phenotypic transitions underlie development, regeneration, and cancer. Our ability to interrogate cell state transitions and their consequences has dramatically increased with advances in single-cell genomics (Rood et al., 2022). We can dissect the timing of key events as cells change state (Qiu, Martin, et al., 2024) and identify transient or intermediate states (MacLean, Hong, et al., 2018). Efforts to produce a comprehensive catalogue of cell states are underway (Regev et al., 2017), yet large gaps in our understanding remain: both regarding cell states and even more regarding the transitions they undertake. We do not have satisfactory explanations of what are the initiating factors of a cell state transition, nor what is the relationship between the dynamics of cell phenotypic change and the transcriptional dynamics acting within the cell.

The epithelial-to-mesenchymal transition (EMT), during which epithelial cells become mesenchymal or mesenchymal-like (Thiery, 2003), is an exemplary cell state transition. EMT is necessary during development and wound healing and is co-opted by cancer, where it is a crucial component of metastasis. Understanding EMT is thus imperative to slowing or preventing metastasis, the leading cause of death from cancer (Dillekås et al., 2019). Classical conceptions of EMT characterize a binary process, with cells being either completely epithelial or mesenchymal (Thiery, 2003). However, experimental and theoretical studies have demonstrated the existence of EMT intermediate states (Bracken et al., 2008; Deshmukh et al., 2021; Hong, Watanabe, et al., 2015; Hong and Xing, 2024; Sha, Haensel, et al., 2019; Xing et al., 2019). Pan-cancer studies of intermediate EMT states have revealed insight into transcriptomic signatures underlying EMT transformation (Tagliazucchi et al., 2023). The intermediate state displays partial EMT phenotypes, with characteristics of both epithelial and mesenchymal states, and may also be called partial EMT, hybrid EMT, or an E/M state (Nieto et al., 2016). EMT intermediate states are closely tied with the concept of epithelial-mesenchymal plasticity (EMP): dynamic, bidirectional transitions through multiple EMT states.

EMT intermediate states are found in both non-malignant EMT and cancer (Nieto et al., 2016; Shaw et al., 2016). The relevance of targeting these states is compelling: EMT intermediate states have been associated with circulating tumor cells (Ruscetti et al., 2015; Ting et al., 2014; Yu et al., 2013) and metastasis (Hendrix et al., 1997), perhaps even more potently than mesenchymal cells alone (Jolly et al., 2015; Simeonov et al., 2021). We focus here on stable EMT intermediate states: biologically, this refers to cells in a state that can be isolated and persist under sufficient conditions; mathematically, stability is defined via the Lyapunov exponents of a dynamical system (Jost et al., 2015). EMT intermediate states have been described as “metastable” in the literature, which in this case refers to stable cell states with small basins of attraction. EMT intermediate state cells may be hard to observe in part due to their rarity (small population sizes or small basins of attraction) or their location (existing at the margins rather than throughout a tissue [Leggett et al., 2021]), although they are not necessarily a minority of cells in a sample.

Mathematical models of EMT have predicted and identified intermediate states, using transcriptional networks that can successfully capture both the steady states of the system and its dynamic properties (Chaffer et al., 2016; Hong, Watanabe, et al., 2015; Lu et al., 2013; Medici et al., 2008; Tian et al., 2013; Zhang, Tian, et al., 2014). These transcription models of EMT, typically regulated by transforming growth factor-beta (TGF-*β*), primarily focus on a core network with transcription factors ZEB, SNAIL, and OVOL, and micro-RNAs miR200 and miR34. Although greater attention has been paid to the transcriptional dynamics, there has also been mathematical modeling of the cell population dynamics during EMT (reviewed in Tripathi et al., 2021).

Integrating single-cell genomics with mathematical models offers means to infer dynamic properties from high-dimensional systems (Cho et al., 2022; Wu et al., 2023). EMT, with its relatively straightforward trajectory (non-branching, non-cyclical), lends itself well to analysis via trajectory inference (pseudotime) (Saelens et al., 2019), albeit not taking into account the spatial components of the cell fate decisions which can be decidedly more complex (MacLean, Smith, et al., 2017). Trajectory inference coupled with mathematical modeling has led to insight into the initiation and timing of EMT (Sha, Wang, Zhou, et al., 2020). Despite limitations in inferring Markovian cell dynamics from single-cell data (Weinreb et al., 2018), experimental methodology such as metabolic labeling (Qiu, Zhang, et al., 2022) or lineage tracing (Simeonov et al., 2021) can overcome these challenges. Here we take an alternative approach to inferring the population dynamics model directly from data (Fischer et al., 2019; Weinreb et al., 2018), and (in keeping with the observation that cell state transition dynamics are non-Markovian [Stumpf et al., 2017]) we propose a population model of EMT cell state transitions *a priori*. We subsequently learn rates of cell state transition for each individual sample via Bayesian parameter inference of the cell dynamics over pseudotime.

Here we use single-cell RNA sequencing (scRNA-seq) data to fit mathematical models of EMT population dynamics across various tumor types and stimuli. Parameter inference across these different conditions reveals shared and distinct properties of the routes of EMT. We identify shared genes associated with EMT intermediate states across tumor types via differential expression and differential RNA velocity analyses. By comparing intermediate state genes with inferred EMT parameters, we identify genes associated with EMT dynamics – that is, genes that speed up or slow down EMT. We confirm top predictions by an independent analysis of EMT in a new cell type, demonstrating how these methods offer novel means to identify biomarkers or potential targets during cell state transitions.

## 2 METHODS

### 2.1 Single-cell data acquisition and processing

#### 2.1.1 Data sources

In this study, we conducted an integrated analysis of several single-cell RNA sequencing (scRNA-seq) datasets in the public domain. We included datasets from Pastushenko et al., 2018 (GEO accession GSE110357); van Dijk et al., 2018 (GSE114397); Cook and Vanderhyden, 2020 (GSE147405); and Panchy et al., 2022 (GSE213753). For data from Cook and Vanderhyden (2020), samples collected after the removal of the EMT stimulus were not included. For data from Panchy et al. (2022), unstimulated cells were not included.

#### 2.1.2 Sequence alignment

Data from Cook and Vanderhyden (2020) were re-aligned to obtain spliced and unspliced read counts for RNA velocity analysis below. Raw sequence files (accession SRP253729) were downloaded from the NIH Sequence Read Archive using the SRA Toolkit (Leinonen et al., 2011) and converted from SRA to FASTQ files using fasterq-dump. Python package cutadapt was used to trim the barcode sequences to 26 base pairs (Martin, 2011). The splici (spliced+intron) index was constructed using the GRCh38 human reference genome with Python package salmon (Srivastava et al., 2019). Sequence pseudoalignment was performed with salmon alevin-fry. Barcode demultiplexing was carried out using the R package MULTIseq (McGinnis et al., 2019). Contaminant cells in the OVCA420 samples were removed as noted by the original authors.

#### 2.1.3 scRNA-seq data preprocessing and normalization

All scRNA-seq data were processed and analyzed using Scanpy (Wolf et al., 2018). Cells with fewer than 200 genes and genes expressed in fewer than three cells were filtered out. Cells with high mitochondrial percentages or disproportionately high total read counts were excluded based on dataset-specific cutoffs (Supp. Table 1). In HMLE samples stimulated with TGF-*β*, cells with disproportionately high ribosomal percentages were filtered out (<1% of cells). Counts were normalized to 10,000 and *log*(*x* + 1) transformed. Batch correction for samples from Cook and Vanderhyden (2020) was performed using ComBat in Scanpy (Johnson et al., 2007). Cell cycle effects, which significantly impacted clustering by EMT state identity, were regressed out (Satija et al., 2015; Tirosh et al., 2016), similar to the original analyses. Additional preprocessing included regressing out total counts and percent mitochondrial counts per cell, scaling counts to uniform variance, and selecting highly variable genes for downstream analysis.

#### 2.1.4 Cell clustering and scoring by EMT status

Principal component analysis (PCA) was performed, and the top 15 components were used to construct a nearest neighbor graph. Based on this graph, cell clustering was conducted using the Leiden algorithm (Traag et al., 2019) with dataset-specific resolutions (Supp. Table 1). Differentially expressed genes for each cell cluster were identified using Wilcoxon rank-sum test with Benjamini-Hochberg correction. Cell clusters were visualized in two dimensions using UMAP and PHATE (McInnes et al., 2020; Moon et al., 2019).

To infer the EMT status of single cells based on a set of EMT marker genes, an “EMTscore” was created using the UCell scoring method (Andreatta et al., 2021) with the Hallmark EMT gene set from the Molecular Signatures Database (MSigDB) (Liberzon et al., 2015; Subramanian et al., 2005). UCell calculates single-cell gene expression scores from a gene set using a rank-based approach, which we found to effectively quantify EMT across disparate tumor types and experimental conditions. Genes were input into UCell as filtered and normalized counts.

#### 2.1.5 Identifying shared EMT intermediate state genes

Genes were included in the intermediate state analysis if they were differentially expressed (DE) in an intermediate state with a Benjamini-Hochberg adjusted Wilcoxon rank-sum p-value of *p <* 0.01, up to a maximum of 500 genes per sample. To account for the complexities of comparing gene expression across different datasets and conditions (e.g. batch effects, instrumentation, sequencing depth), we calculated log_2_ fold change (log_2_FC) values of intermediate state genes ins Scanpy, i.e. according to (following the notation of (Moses et al., 2023)):

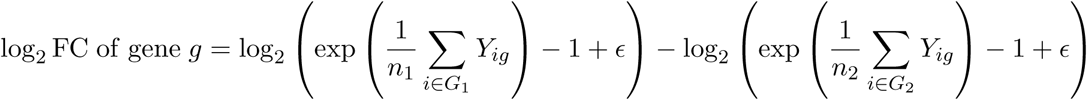

where *G*_1_ is the focal group of cells of size *n*_1_ cells, *G*_2_ is the comparison group with *n*_2_ cells, and *Y_ig_* denotes the log-normalized counts of gene *g* in cell *i*. The pseudocount *ε* = 10*^−^*^9^ is added to avoid division by zero (Moses et al., 2023). Genes were selected as intermediate state-associated if they met the following criteria: i) a log_2_FC *≥* 0.58 (1.5-fold change) in at least five samples, and ii) at least two of these samples were from experiments not performed on HMLE cells.

### 2.2 Trajectory inference & EMT subpopulation dynamics

Diffusion pseudotime (DPT) was used for trajectory analysis (Haghverdi et al., 2016). Root nodes were chosen as the epithelial cells with extreme coordinates on a diffusion map. Pseudotime was calculated five times with different epithelial root nodes, and the median values were assigned to each cell, with the standard deviation indicating pseudotime variation. This approach minimized the impact of root node selection on pseudotime calculation. Pseudotime values range from 0 (epithelial) to 1 (mesenchymal). This range was divided into fifteen bins (twelve for Pastushenko et al. (2018) due to fewer cells) and cell counts were calculated for each cluster (epithelial, intermediate, and mesenchymal) for each bin. The counts per bin were converted into cell population proportions.

### 2.3 RNA velocity analysis

RNA velocity analysis was conducted in Python using the package scVelo in dynamical mode on highly variable genes (Bergen et al., 2020). Each sample was analyzed individually. Differential velocity (DV) was assessed using the rank_dynamical_genes function on clusters. Genes with a DV score above 0.25 were retained as DV genes. To ensure monotonic transitions, genes with Spearman correlation coefficients below 0.5 were excluded. Additionally, DV genes with poor dynamical model fits were filtered out. Ultimately, we retained DV genes that were upregulated in the majority of cancer samples, designating them as shared upregulated velocity genes across EMT.

### 2.4 A mathematical model of EMT dynamics

We developed a mathematical model of the dynamics of EMT described by ordinary differential equations (ODEs). Specifically, we sought to describe the cell state transitions during EMT, from the epithelial (*E*) to intermediate (*I*) state or states, and then to the mesenchymal (*M*) state. While EMT systems may also exhibit direct transitions (*E → M*) and reverse transitions, our data specifically investigate forward EMT and do not exhibit strong evidence for direct transitions.

The population dynamics of *E*, *I*, and *M* are described by:

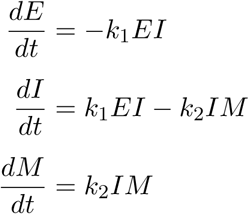

where *k*_1_ denotes the transition rate from *E* to *I*, and *k*_2_ denotes the transition rate from *I* to *M*. We consider second-order transitions, meaning both the initial and final states influence the transition rate to the final state. In cases where more than one intermediate state exists, the model can be extended using the same framework (Supp. Fig. 7).

### 2.5 Parameter inference of cell population dynamics over pseudotime

We sought to infer the rates of EMT using Bayesian parameter inference with the Turing.jl package in Julia (Bezanson et al., 2017; Ge et al., 2018; Rackauckas et al., 2017). The input data for each model consists of the cell state dynamics over pseudotime. To focus on relevant dynamics, we excluded periods where all cells remained in the epithelial state. Time points along pseudotime were normalized to a range of *t ∈* [0, 10], facilitating direct comparison of EMT trajectories across samples. For each sample with one intermediate state, we fit three parameters: *k*_1_, *k*_2_, and the observational noise parameter *σ*. For the *in vivo* sample with two intermediate states, we fit four parameters: *k*_1_, *k*_2_, *k*_3_, and *σ*.

Letting *f* represent the numerical solution to the ODE model and *y*_0_ the initial conditions, we performed parameter inference as follows:

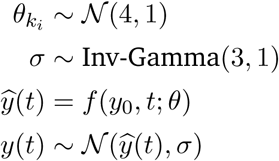

where *θ* = (*θ_k_i__, σ*) gives the prior parameter distribution, and *y*(*t*) defines the likelihood function in terms of ODE model simulation (*ŷ*(*t*)) for transition rate parameters *θ_k_i__* and noise parameter *σ*.

The posterior parameter distribution was estimated via Markov chain Monte Carlo (MCMC) simulations using the No-U-Turn Sampler (NUTS) (Hoffman et al., 2014). MCMC chains were each run for 1,000 iterations following 250 warmup iterations to ensure convergence. Fitted trajectories were visualized by solving the model using 300 joint parameter sets of *k_n_*, randomly selected from the posterior distribution for each sample, and plotting the mean and standard deviation of the resulting trajectories.

### 2.6 Comparative analysis of EMT intermediate state-associated genes

We identified genes associated with EMT transition rates by analyzing correlations between model-inferred posterior parameters and gene expression. For each transition rate parameter *k_n_*, we used its maximum *a posteriori* value for each sample and examined pairwise correlations with the log_2_FC expression of 145 genes, each present in at least 5 samples with an intermediate state log_2_FC *≥* 0.2. Genes with a Spearman’s rank correlation coefficient of *ρ > |*0.6*|* (*p <* 0.05) were considered associated with, and potential influencers of, transitions into or out of EMT states.

To specifically identify genes linked to EMT intermediate state dynamics, we focused on genes positively correlated with *k*_1_ (faster *E → I*) and negatively correlated with *k*_2_ (slower *I → M*). Genes meeting both correlation criteria were included, as well as those showing either correlation as well as differential expression or differential velocity in the intermediate state. Cellular location annotations were performed using DAVID (Huang et al., 2009; Sherman et al., 2022) and PANTHER (Thomas et al., 2022).

## 3 RESULTS

### 3.1 Single-cell analysis of EMT across cancer types & stimuli identifies a spectrum of EMT states

To characterize trajectories across a spectrum of EMT, we studied twelve scRNA-seq datasets across five cancer types. Cells were processed and clustered to identify cell states. We found evidence for three cell states in each of the *in vitro* cell populations and four states in the *in vivo* mouse skin squamous cell carcinoma (SCC) sample (Supp. Fig. 1 & 2). Clusters were labeled based on EMT markers from the literature, including Hallmark EMT genes from the Molecular Signatures Database (MSigDB) (Liberzon et al., 2015) and epithelial cell genes from PanglaoDB (Franzén et al., 2019). Distinct clusters representing epithelial and mesenchymal cell types were identified in each dataset, although the relative sizes of these clusters varied widely (Fig. 1A). In all datasets, at least one cluster expressing combinations of epithelial and mesenchymal marker genes was identified as an intermediate state. Certain samples from Cook and Vanderhyden (2020) that did not exhibit a clear EMT were excluded from further analyses (Supp. Fig. 3 & 4). This is in agreement with Cook and Vanderhyden (2020), who also found that certain conditions did not permit a full EMT within the experimental timeframe.

**Figure 1:**
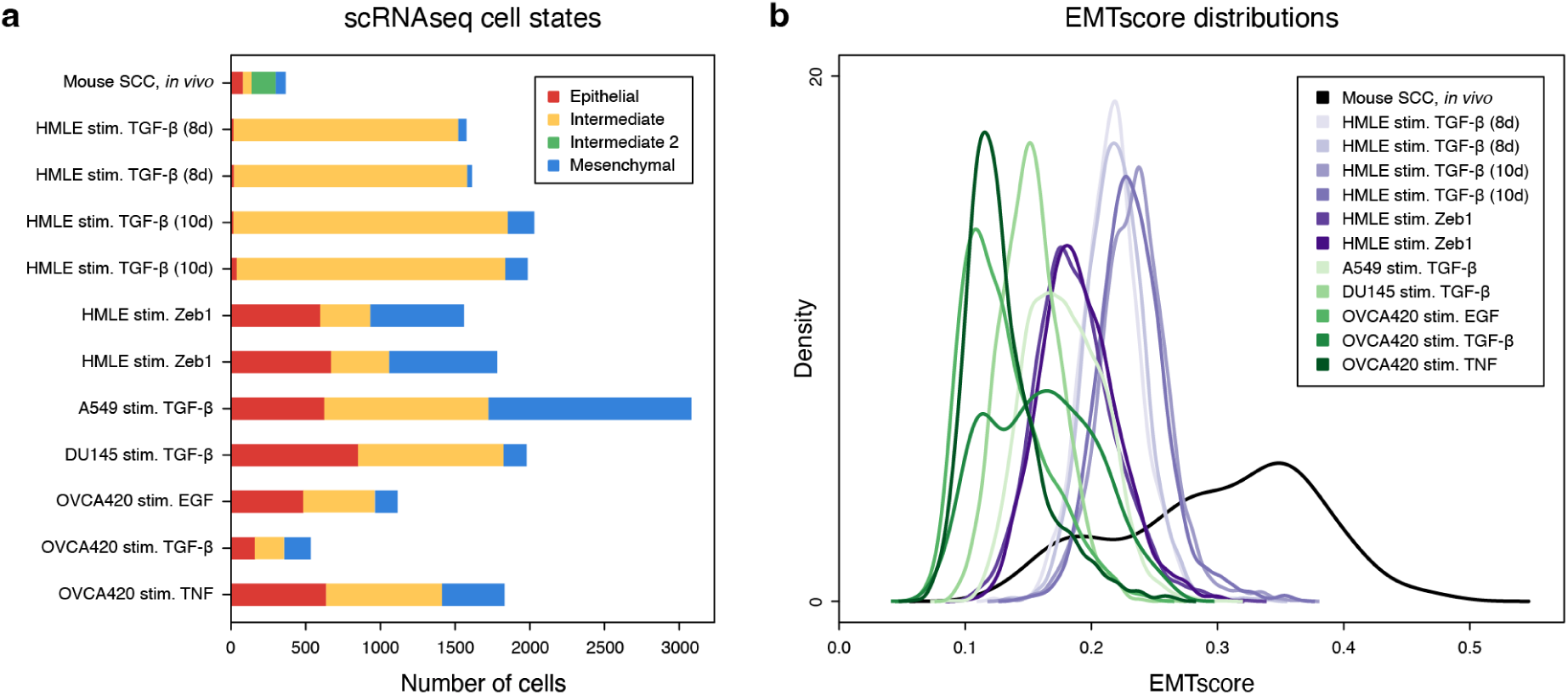
**a.** Cell counts of EMT states per cancer sample, including intermediate states. Cell states were identified via clustering and gene expression. **b.** Kernel density estimates of the EMTscore distributions for each scRNAseq sample. Single-cell EMT scores were assigned via Hallmark EMT genes from MSigDB.

EMT scores were assigned to single cells across all datasets (Fig. 1B). Single cells were each assigned an EMTscore via UCell (Andreatta et al., 2021) using MSigDB Hallmark EMT genes. Each sample exhibited a range of EMTscore, reproducible by replicate and varying considerably by cell type and stimulus. Notably, not only the variance but also the start and end points vary by cell type, highlighting differences not only in EMT but also in the “epithelialness” of different cell types. Samples excluded from analysis due to lack of/incomplete EMT, as identified by marker gene expression, exhibited little to no variation in EMTscore (Supp. Fig. 4), confirming the lack of cell state transition under the tested conditions. The *in vivo* EMT in mouse SCC exhibited the largest range of EMTscore by a wide margin, highlighting the increase in heterogeneity among single cells during a spontaneous, unstimulated, environment-dependent EMT. Since an additional intermediate state was identified in this dataset, in line with previous work (Pastushenko et al., 2018; Sha, Wang, Bocci, et al., 2021), the data suggest that both the number of attractor states and the size of their basins of attraction are larger for cells in their natural environment than cell line-derived models stimulated *in vitro*.

### 3.2 Shared marker genes of intermediate EMT states are associated with extracellular function

EMT can proceed along many paths (Hong and Xing, 2024), and both cell/treatment-specific and consensus EMT pathways are important to study in different contexts. Here, we focus on the shared properties of EMT cell state transitions. To study intermediate state gene expression across an EMT spectrum, we performed differential gene expression and differential RNA velocity analysis across intermediate states in different cell populations (Fig. 2A). We identified differentially expressed genes for intermediate states in each sample (2,396 genes total) and examined shared intermediate state-specific genes, defined as those upregulated in an intermediate state relative to epithelial/mesenchymal states. Using a log_2_ fold change (log_2_FC) threshold of +0.58 (1.5-fold change) in at least five samples, we identified 32 genes shared among EMT intermediate states (Supp. Fig. 5).

**Figure 2:**
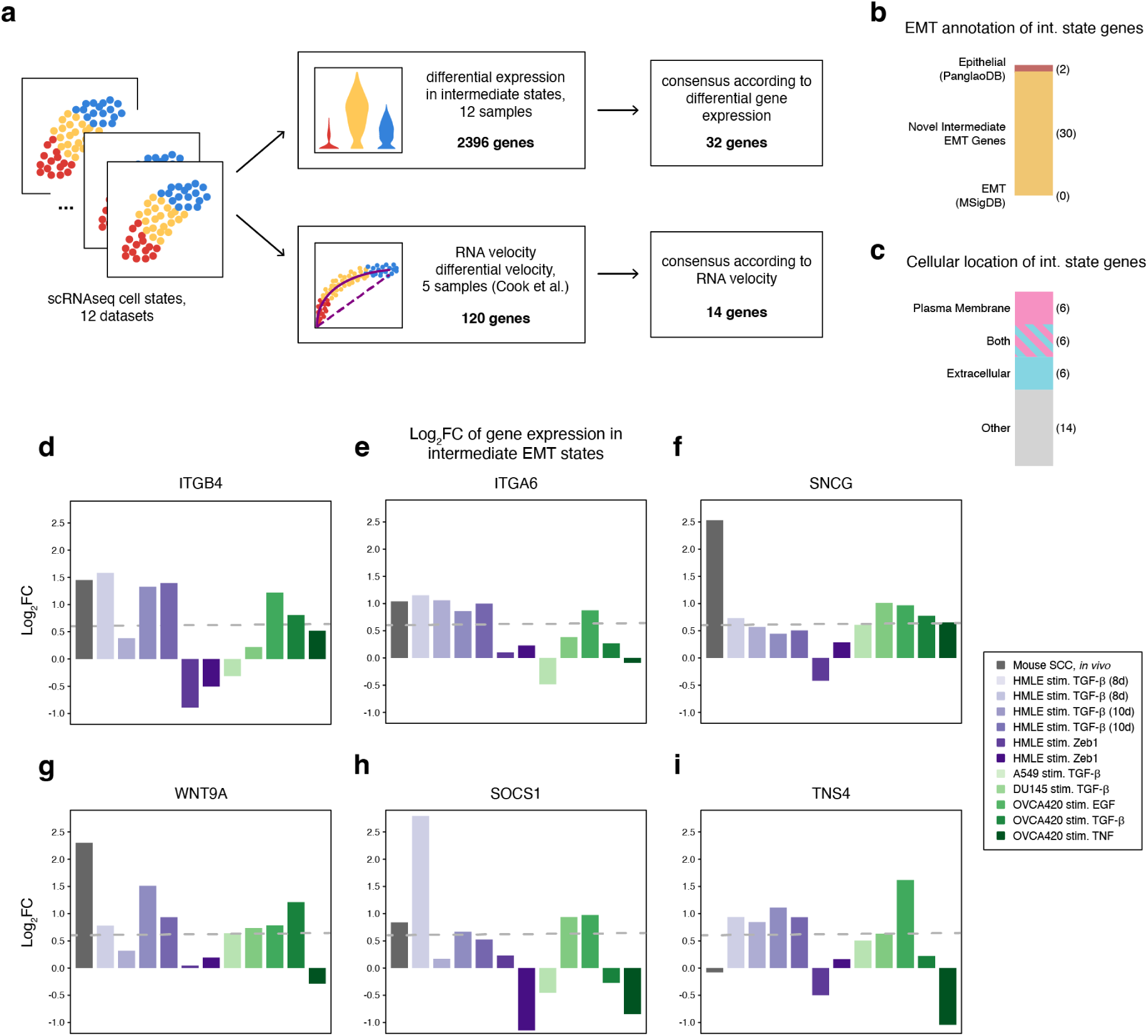
**a.** Data analysis pipeline: pan-cancer intermediate EMT marker genes were identified using differential expression and compared against genes differentially regulated via RNA velocity. **b.** Annotation of predicted intermediate EMT genes by canonical epithelial/mesenchymal gene sets. **c.** Annotation of predicted intermediate EMT genes by cellular location. **d-i.** Comparison of expression of predicted intermediate EMT genes by log_2_FC (in the intermediate state) across samples. Dashed line represents 1.5-fold change (log_2_FC > 0.58).

Among the 32 genes shared across EMT intermediate states, most were not found in canonical EMT or epithelial gene sets (Fig. 2B). Two predicted intermediate state genes, *ITGB4* and *SFN*, are annotated as epithelial genes in PanglaoDB (Franzén et al., 2019), although the literature on these genes is complicated: Integrin *β*4 (*ITGB4*) (Fig. 2D) was initially identified in epithelial cells and tumors (Biffo et al., 1997) but has also been linked to promoting EMT in hepatocellular and pancreatic carcinoma (Li et al., 2017; Masugi et al., 2015). *ITGB4* pairs with another intermediate state gene, integrin *α*6 (*ITGA6*) (Fig. 2E), to form the *α*6*β*4 complex, which is implicated in promoting EMT characteristics in hepatocellular carcinoma cells (Zheng et al., 2022). Stratifin (*SFN*) (Supp. Fig. 5) is annotated as epithelial (named for its role in the stratification of epithelial cells [Leffers et al., 1993]) but is also linked to cell migration and EMT markers in cervical and hepatocellular carcinoma (Hu et al., 2019; Ye et al., 2023; Zhao et al., 2023). The apparent contradictory roles of both *ITGB4* and *SFN* as marking for both epithelial and mesenchymal states can be reconciled if these genes are in fact markers of an intermediate EMT state, as predicted by our analysis.

A majority of predicted intermediate EMT marker genes encode proteins localized in the extracellular space, on the plasma membrane or as secreted signaling factors (Fig. 2C). Gamma-synuclein (*SNCG*), upregulated across multiple cell lines (Fig. 2F), is found in the extracellular exosome. It plays a role in suppressing mesenchymal markers including *CDH2* (N-cadherin) and *VIM* (Ni et al., 2021) while promoting cancer cell migration (Liu et al., 2022; Takemura et al., 2021; Zhuang et al., 2015). Other notable upregulated genes include *WNT9A*, *IL4R*, and *IL6R*. Wnt-9a (*WNT9A*) (Fig. 2G) is a secreted protein in the canonical Wnt/*β*-catenin signaling pathway that is implicated in partial EMT by mediating cell adhesion (Basu et al., 2018). *IL4R* and *IL6R* (Supp. Fig. 5) are interleukin cell surface receptors, with their cytokines IL4, IL13, and IL6 associated with EMT promotion (Cao et al., 2016; Chen et al., 2018; Sun et al., 2020). Interestingly, *SOCS1* (suppressor of cytokine signaling 1) (Fig. 2H) is a negative regulator of IL6 yet conversely has been found to promote EMT (Berzaghi et al., 2017), highlighting the bidirectional signaling at play during the establishment of intermediate EMT states.

Tensin 4 (*TNS4*) (Fig. 2I) is involved in focal adhesion & integrin interaction and promotes EMT and cell motility (Katz et al., 2007; Thorpe et al., 2017). Tubulointerstitial nephritis antigen-like 1 (*TINAGL1*) (Supp. Fig. 5) encodes another secreted protein that binds directly to certain integrins, and it is found to both promote and inhibit metastasis in different cancers *in vivo* (Shan et al., 2021; Shen et al., 2019). Both *TNS4* and *TINAGL1* interact with epidermal growth factor receptor *EGFR*, yet their effects are contradictory: *TNS4* reduces *EGFR* degradation (Hong, Shih, et al., 2013), while *TINAGL1* binds directly to *EGFR* and suppresses *EGFR* signalling (Shen et al., 2019). These opposing interactions may again reflect the dynamic balance necessary to sustain the intermediate EMT state.

Overall, many genes associated with the intermediate EMT state exhibit conflicting roles in the literature, including *ITGB4*, *SFN*, *IL4R* & *IL6R* with *SOCS1*, and *TINAGL1* with *TNS4*. These genes can contribute both to the promotion and inhibition of EMT as well as the balance between epithelial and mesenchymal states. This duality underscores the dynamic nature of EMT and the importance of intermediate states.

### 3.3 Differential regulation via RNA velocity reveals EM plasticity genes in EMT intermediate states

To investigate dynamically regulated genes during EMT, we performed differential RNA velocity across EMT cell states (Bergen et al., 2020; La Manno et al., 2018). Fourteen genes had differential velocity (DV) in the intermediate state in at least three of the five Cook and Vanderhyden (2020) samples, which includes cells from lung, prostate, and ovarian tumors (Supp. Fig. 6). Of the 14 DV genes, all but one encode proteins located extracellularly or in the plasma membrane (Fig. 3A). Several of these genes are involved in focal adhesion, including integrins *ITGA2* and *ITGB4*, laminins *LAMC2* and *LAMB3*, collagen *COL4A2*, and plasma membrane caveolae component *CAV1*. Eleven of the DV genes have annotated signal peptide sequences, underscoring their designation as secretory/membrane proteins (Sherman et al., 2022; The UniProt Consortium, 2023).

**Figure 3:**
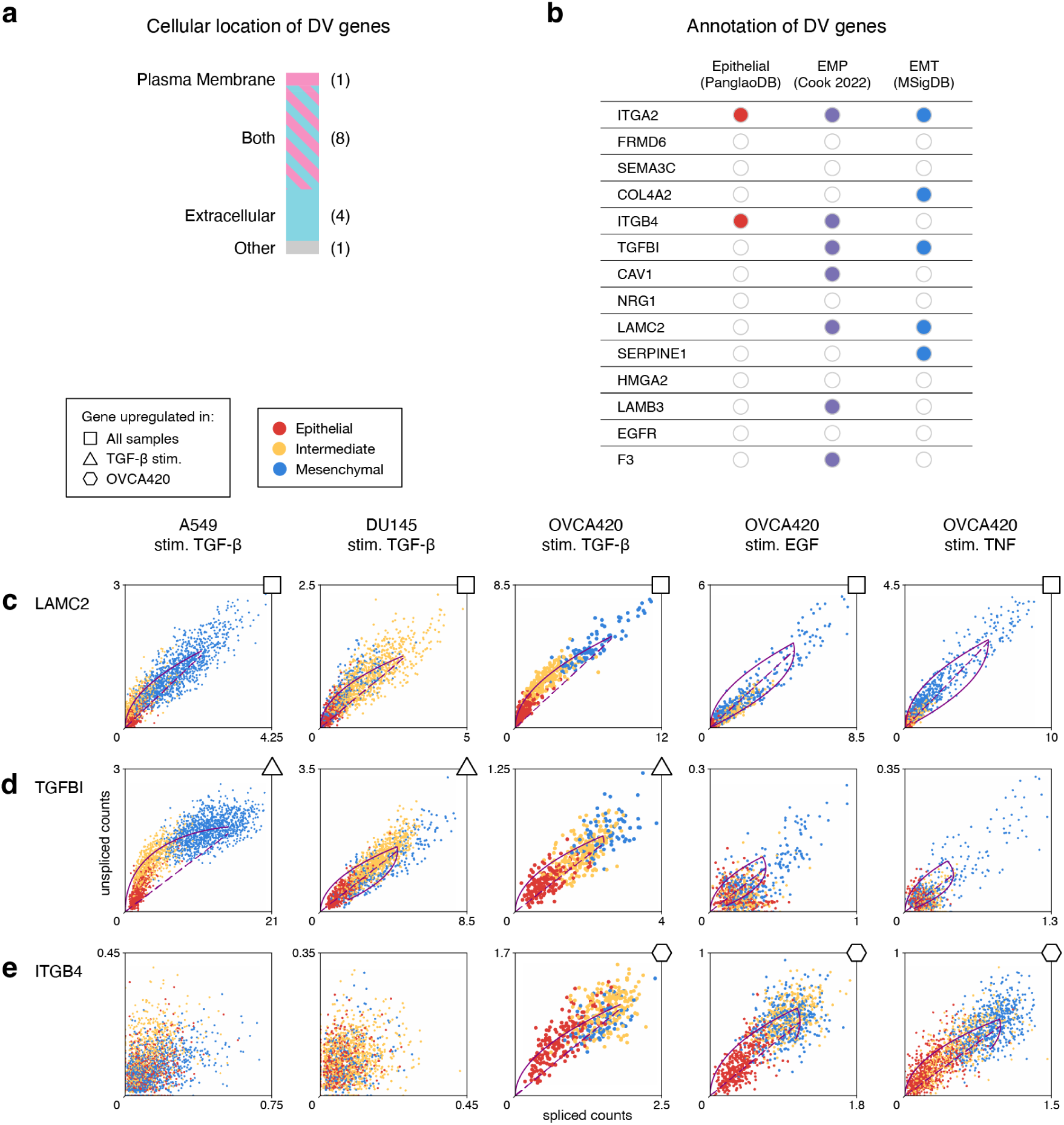
**a.** Annotation of cellular locations for genes differentially regulated in the intermediate state, identified through differential velocity (DV) analysis. **b.** Annotation of the EMT properties of DV genes by comparison with three EMT marker gene sources. EMP: epithelial-mesenchymal plasticity. **c.** Examples of DV genes upregulated in intermediate EMT states across different conditions. Solid line represents the dynamical model fit; dashed line represents the inferred steady state. LAMC2 is upregulated across samples from different cell lines & stimuli. **d.** ITGB4 is upregulated only with a specific stimulus: TGF-β. **e.** ITGB4 is upregulated only in a specific tumor type: ovarian cells OVCA420.

DV genes showed greater overlap with canonical EMT gene sets than the intermediate state marker genes we identified (Fig. 3B). This is expected, as genes actively upregulated in EMT intermediate states are more likely to overlap with mesenchymal markers. Epithelial-mesenchymal plasticity (EMP), i.e. bidirectional cell state transitions between epithelial and mesenchymal phenotypes (Cook and Vanderhyden, 2022), is also characteristic of the DV genes identified. This overlap supports EMP conceptually: capricious cells require dynamic changes in gene expression to change state.

Comparison of DV genes across samples revealed a variety of responses: some genes were shared across different cell types and conditions, while others were specific to certain conditions. Genes upregulated regardless of cell type or stimuli included *LAMC2* (Fig. 3C), *FRMD6*, and *SERPINE1* (Supp. Fig. 6). In contrast, and perhaps unsurprisingly, TGF-*β*-induced protein *TGFBI* (Fig. 3D) was upregulated in various cell types only when stimulated by TGF-*β*. A similar pattern was observed for *COL4A2* (Supp. Fig. 6). Genes upregulated by multiple stimuli in one cell type, human ovarian OVCA420 cells, included *ITGB4* (Fig. 3E), *CAV1*, *HMGA2*, *F3*, and *LAMB3* (Supp. Fig. 6). Overall, RNA velocity analysis elucidates gene regulation during EMT. Most differentially regulated genes are specific to a stimuli or cell line, fewer are conserved across conditions. There is substantial overlap between actively regulated genes during EMT and those linked to EMP, highlighting the role of dynamic transitions between cell states during EMT.

### 3.4 Mathematical modeling & parameter inference quantifies EMT population dynamics

Gene expression is not static: life arises from dynamics. To study the dynamics of EMT in more depth, we developed a mathematical model describing cell state transitions during EMT (Fig. 4A). The model is characterized by rate parameters for transitions between epithelial (*E*), intermediate (*I*), and mesenchymal (*M*) states, such as *E → I* at rate *k*_1_ (Fig. 4B; Supp. Fig. 7A). These rate parameters were fit to scRNA-seq data, characterizing cell state transitions during EMT across pseudotime. Multiple pseudotime trajectories were calculated for each sample, rooted by different epithelial cells, to estimate the mean & variance in pseudotime based on root node selection. Cell state proportions across pseudotime, representing cell population dynamics during EMT, were fitted to the model.

**Figure 4:**
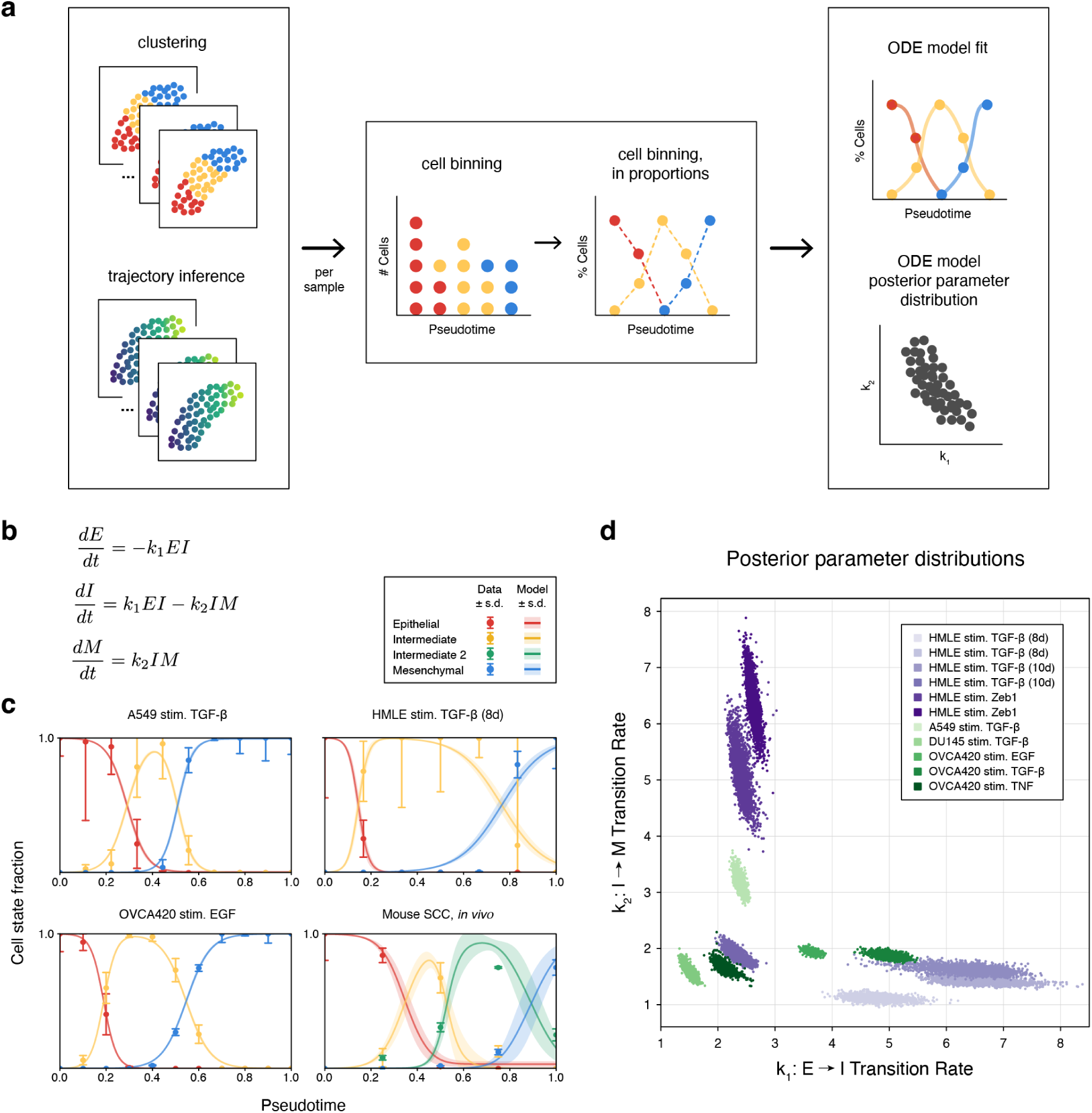
**a.** Workflow to infer dynamic EMT transition rates from scRNA-seq data. For each sample, clustering and trajectory inference information was processed to quantify cell states over pseudotime. A mathematical model was then fit to each sample to infer parameter posterior distributions. **b.** Mathematical model representing transitions from epithelial (E) to intermediate (I) to mesenchymal (M) state cells. k_1_ is the transition rate E → I; k_2_ is the transition rate I → M. Additional intermediate states can be seamlessly added (Supp. Fig. 7A). **c.** Model fits following parameter inference: data vs. trajectory simulations, with simulation parameters sampled from the posterior of each model. **d.** Posterior parameter distributions of the model for each fitted sample.

EMT dynamics for each dataset were fit using Bayesian parameter inference (Fig. 4C; Supp. Fig. 7; Supp. Table 2). Differences in EMT dynamics were observed across different datasets, both by cell type and by stimulus. For instance, the intermediate state persisted longer in HMLE cells compared to A549 or OVCA420 cells. Analysis of the parameter posterior distributions for each fitted EMT trajectory revealed similarities and differences in EMT dynamics (Fig. 4D). Dividing the posterior space into three approximate regions: *k*_1_ *≈ k*_2_ (similar transition rates across EMT); *k*_1_ *> k*_2_ (faster transition rates for *E → I* than *I → M*); and *k*_1_ *< k*_2_ (faster transition rates for *I → M* than *E → I*) highlights how both cell type and stimulus can strongly impact EMT dynamics. For example, OVCA420 cells exhibited *k*_1_ *> k*_2_ dynamics regardless of stimulus, where *k*_1_ *> k*_2_ implies a larger/more stable intermediate state. In contrast, HMLE cells exhibited *k*_1_ *> k*_2_ dynamics for TGF-*β* stimulation but *k*_1_ *< k*_2_ for ZEB1 stimulation, indicating that the persistence/stability of the HMLE intermediate state depends on the stimulating factor. An inverse proportion relationship is evident across cell types/stimuli and within a sample; this concordance is notable since more generally different types of parameter covariation can exist (Wu et al., 2023). This analysis highlights how EMT intermediate persistence and stability depend on the intrinsic properties of the EMT experiment, with different carcinomas exhibiting greater or lesser sensitivity to EMT-inducing factors and thus affecting EMT progression.

### 3.5 Consensus analysis predicts that *SFN* and *NRG1* influence intermediate cell state dynamics during EMT

To identify genes influencing intermediate EMT dynamics, we studied associations between intermediate EMT genes and fitted parameters of the mathematical model. A gene’s positive correlation with *k*_1_ indicates faster transition *E → I*, while a negative correlation with *k*_2_ means a slower transition *I → M*; either correlation suggests that the gene is associated with a more persistent intermediate state. Genes with significant Spearman’s correlation were compared with differential expression and differential velocity genes in intermediate states, and those supported by multiple lines of evidence were consolidated into a consensus gene list of 14 genes (Supp. Table 3; Supp. Fig. 8). The majority of intermediate EMT dynamics genes were located at the plasma membrane or in the extracellular region (Fig. 5A). Of the 14 predicted intermediate EMT dynamics genes, three were identified in a prior EMP study (Cook and Vanderhyden, 2022) (Fig. 5B), consistent with the conceptual overlap between intermediate EMT dynamics and EMP. Notably, there is no overlap between intermediate EMT dynamics genes and those from hallmark EMT (mesenchymal) genes, demonstrating that our proposed gene set is novel and distinct from previous EMT gene sets.

**Figure 5:**
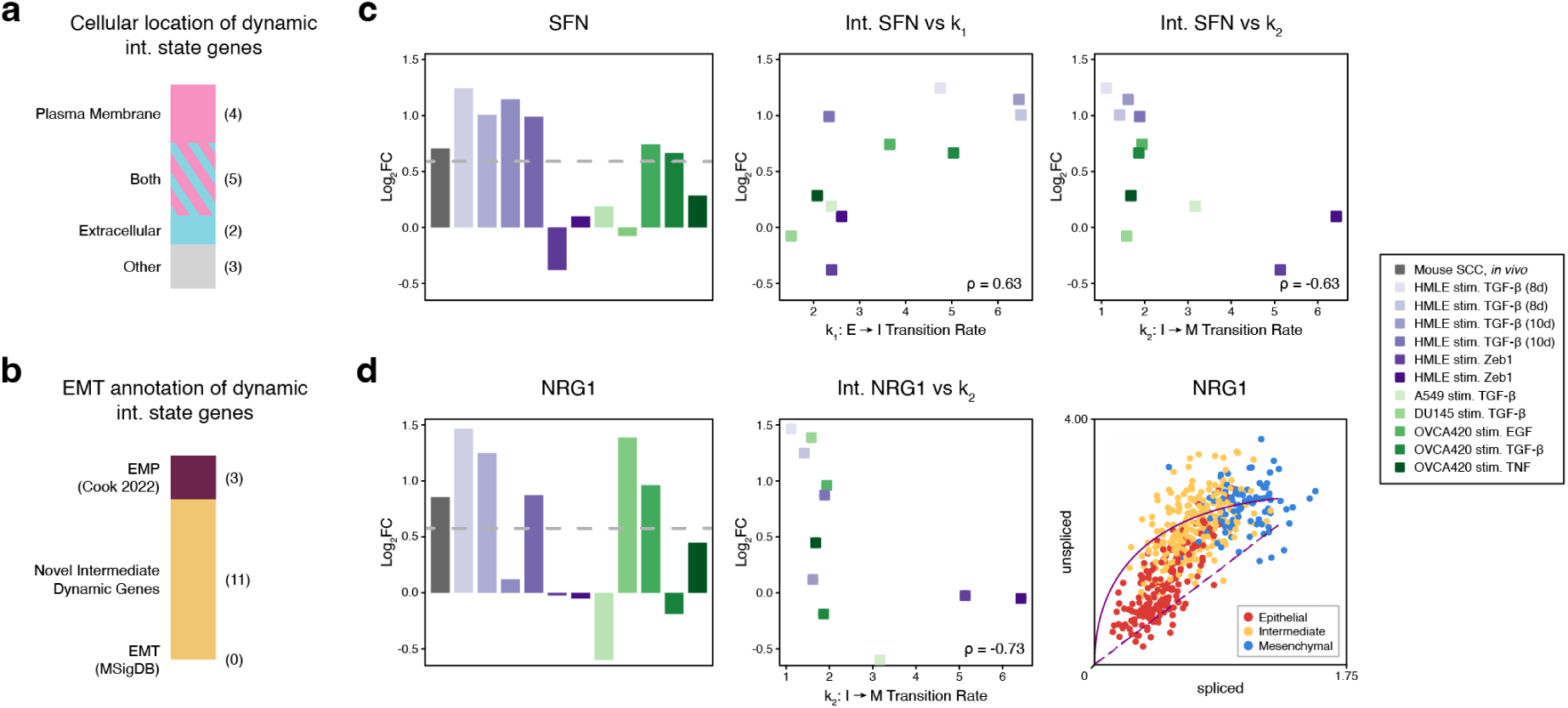
**a.** Annotation of predicted dynamic intermediate state genes by cellular location. **b.** Annotation of predicted dynamic intermediate state genes by EMT marker gene sources. **c.** SFN is a predicted dynamic intermediate state gene, with its differential expression in multiple EMT intermediate states and significant correlations with model parameters k_1_ and k_2_. **d.** NRG1 is another predicted dynamic intermediate state gene, with its differential expression in multiple EMT intermediate states, significant negative correlation with model parameter k_2_, and differential regulation in EMT intermediate states via RNA velocity.

Two predicted EMT dynamics genes had the strongest support (three lines of evidence each; Supp. Table 3). *NRG1* was the only gene identified in all three analyses, while *SFN* was the only gene with intermediate EMT differential expression and significant correlations with both *k*_1_ and *k*_2_ transition rates. Stratifin (*SFN*) was positively correlated with *k*_1_ (*E → I*) and negatively correlated with *k*_2_ (*I → M*) across cancer samples (Fig. 5C), suggesting that it stabilizes the intermediate EMT state. Although RNA velocity for *SFN* was not captured due to insufficient counts, it was differentially expressed in intermediate states. Neuregulin 1 (*NRG1*) was negatively correlated with *k*_2_ (*I → M*), suggesting it slows the exit from the intermediate state (Fig. 5D), and *NRG1* was also significant in intermediate EMT differential expression and velocity (Supp. Fig. 6).

Consensus gene analysis predicts that *SFN* promotes transitions from an epithelial state to the metastatic intermediate EMT state. This prediction helps to reconcile literature, which reports both epithelial and pro-EMT roles for *SFN*. Named for its expression in stratified epithelial cells (Leffers et al., 1993), *SFN* can be secreted and is found in extracellular vesicles (Hou et al., 2022). Recombinant *SFN* treatment has been shown to significantly enhance extracellular matrix degradation in human dermal fibroblasts *in vitro* (Ghaffari et al., 2006). Despite its epithelial association, *SFN* knockdown in *in vitro* models has led to reduced mesenchymal marker expression in cervical cancer cells (Hu et al., 2019) and decreased cell migration in other carcinomas (Kim, Kim, et al., 2022; Ye et al., 2023; Zhao et al., 2023). *In vivo*, *SFN* knockdown suppressed tumor formation and metastasis in lung adenocarcinoma models (Shiba-Ishii et al., 2015). Clinically, *SFN* is linked to poor prognosis, including advanced tumor stages in lung adenocarcinoma and hepatocellular carcinoma (Kim, Shiba-Ishii, et al., 2018; Ye et al., 2023), as well as lower survival rates in pancreatic ductal adenocarcinoma (Robin et al., 2020) and head and neck squamous cell carcinoma (Chung et al., 2006). Our findings suggest that *SFN* promotes intermediate EMT dynamics, potentially explaining its dual role in epithelial cells while facilitating EMT.

Consensus gene analysis also identified *NRG1* as playing a pivotal role in intermediate EMT state dynamics, as the sole gene that was significant in intermediate expression, regulation, and modeled dynamics. A member of the epidermal growth factor (EGF) family (The UniProt Consortium, 2023), *NRG1* activates *ERBB2* (*HER2*) and *ERBB3* (*HER3*) (Miano et al., 2022). *NRG1* isoforms can be found in the plasma membrane or secreted (Esper et al., 2006), and it binds integrins including *ITGA6*:*ITGB4* and *ITGAV*:*ITGB3* (Ieguchi et al., 2010). *In vivo*, *NRG1* suppression reduces tumor growth and metastasis in hepatocellular carcinoma (Shi et al., 2018). Clinically, *NRG1* overexpression correlated with poor outcomes, including lymph node metastasis, in gastric cancer (Yun et al., 2018). Notably, *NRG1* has been found to promote partial EMT in cultured patient *HER2*-positive breast cancer (Guardia et al., 2021). While *NRG1* has been mostly described to drive EMT in epithelial cells, *NRG1* stimulation on mesenchymal cells that already underwent EMT has been shown to instead induce epithelial gene expression in esophageal adenocarcinoma (Ebbing et al., 2017). Taken together, our analyses along with literature suggest that *NRG1* is a marker of highly plastic intermediate state cells during EMT.

### 3.6 *SFN* is a marker of intermediate state EMT in independently analyzed MCF10A cells

To assess predicted intermediate EMT genes, we analyzed a dataset of EMT under different experimental conditions and in a different cell line: the dose-dependent TGF-*β* stimulation of MCF10A breast cells (Panchy et al., 2022). Similar to previous analyses, scRNA-seq data was clustered, and canonical markers were used to identify epithelial, intermediate, and mesenchymal states (Fig. 6A). Differential expression by cell state showed strong agreement with our predictions, with 11 of the top 25 intermediate state genes in this sample overlapping with our predicted intermediate EMT genes (Fig. 6B), notably including *SFN*. These results highlight that shared EMT intermediate state features can be found across diverse biological and experimental conditions, with independent evidence corroborating one of the top genes associated with intermediate EMT.

**Figure 6:**
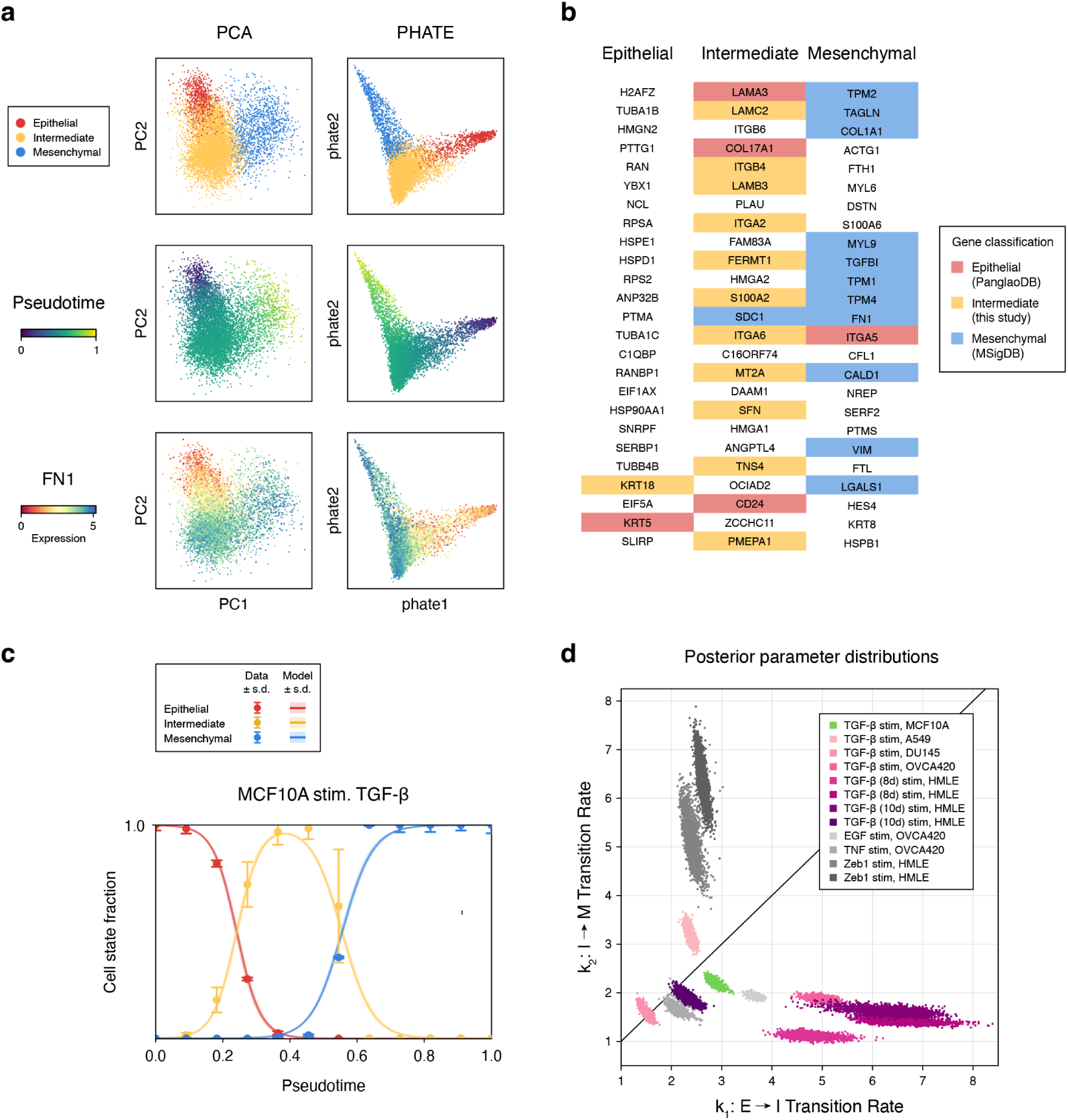
**a.** MCF10A cells were analyzed separately and exhibit a linear trajectory across EMT states. **b.** For each MCF10A cell state, the top 25 differentially expressed genes colored by gene set annotation. **c.** Model fit following parameter inference: data vs. trajectory simulations, with simulation parameters sampled from the posterior distribution. **d.** Comparison of posterior parameter distributions, with the MCF10A sample highlighted in green. Other distributions are replicated from Fig. 4D, shown in different colors here to highlight stimulation by TGF-β.

To further evaluate EMT dynamics in these MCF10A cells, we fit the mathematical model to this dataset (Fig. 6C). The posterior parameter distribution lies in the region where *k*_1_ *≥ k*_2_, consistent with EMT dynamics induced by TGF-*β* in other cell types (Fig. 6D). Across different cancer types, we see that mammary (MCF10A and HMLE) and ovarian (OVCA420) cells stimulated with TGF-*β* generally exhibit *k*_1_ *> k*_2_ dynamics, favoring stabilization of the intermediate state. In contrast, lung (A549) and prostate (DU145) cells stimulated with TGF-*β* show balanced rates of entry and exit from the intermediate state, with *k*_1_ *≈ k*_2_. The similarity in transition dynamics between mammary and ovarian cells is notable, given the shared genetic and microenvironmental factors during oncogenesis and tumor progression (Roskelley et al., 2002).

## 4 DISCUSSION

Here, we characterized intermediate EMT states and identified genes involved in dynamic transitions between states. Multiple lines of evidence suggest EMT intermediate states are the most cancer stem-like and exhibit the highest metastatic potential (Aiello et al., 2018; Brown et al., 2022; Pastushenko et al., 2018; Puram et al., 2017; Zhang, Donaher, et al., 2022). Our analysis predicted intermediate state genes in agreement with recent work, such as *ITGB4* and *LAMB3* (Cheng et al., 2024), as well as novel EMT intermediate genes, such as *SFN* and *NRG1*. While there are many paths of EMT, our comparison across different cell types and stimuli revealed common markers for intermediate states and highlighted the role many of these genes have in extracellular remodeling.

EMT is heterogeneous (Yang et al., 2020). Multiple transcription factors can initiate EMT (Nieto et al., 2016) and act in complex and nonlinear ways, both alone (Hartmann et al., 2024) or in combination (Sanford et al., 2020). Additional factors contributing to EMT complexity, including subtypes and intermediate states, are hysteresis during the reverse mesenchymal-epithelial transition, differences in cell types or stimuli, and state transitions driven by intrinsic or extrinsic noise. Whereas EMT is most frequently modeled via gene regulatory networks, here we modeled the population dynamics to study cell state heterogeneity and its effects on EMT path variation. In doing so, we assumed a monostable landscape, whereas in reality multiple stable steady states exist (Hong, Watanabe, et al., 2015). Some of the gene expression heterogeneity underlying these multiple states is likely collapsed by this approach, but in doing so we can identify consensus genes marking for properties of EMT states across different conditions. Our model can be adapted in the future to consider multiple intermediate states and more complex (e.g. convergent/divergent) EMT paths.

Summarizing complex data across conditions to find consensus requires simplifying assumptions. To compare gene expression across datasets we used log-fold changes and rank-based comparisons, similar to other recent work (Theodoris et al., 2023). Doing so relies on the accurate quantification of cell states, which is not guaranteed, and can obscure single-cell resolution information by taking pseudo-bulk measurements. While we sought to standardize data analysis pipelines as far as possible, scRNA-seq data analysis relies on certain parameter choices. While clustering cells we sought fewer clusters (lower resolution) where supported, to reduce overfitting cell states. Trajectory inference relies on accurate choice of root cells and the sufficiency of the similarity metric used. RNA velocity analysis is limited by the ratio of spliced to unspliced counts, typically around 75-85% spliced to 15–25% unspliced (La Manno et al., 2018). This abundance limitation affects genes with low or no unspliced counts, such as *SFN* in our study, where RNA velocity analysis could not be performed due to a lack of unspliced counts. This abundance limitation could be addressed by experimental methods targeting dynamics, such as RNA metabolic labeling (Qiu, Zhang, et al., 2022).

Mathematical modeling and parameter inference with single-cell data allow us to investigate the genes and pathways associated with dynamic transitions between states rather than the cell states themselves – transitions which are strongly relevant to epithelial-mesenchymal plasticity (Williams et al., 2019). EMP, exemplary of cell state plasticity, has been shown to play decisive roles in tumorigenesis and cancer progression (Househam et al., 2022; Street et al., 2023). This property can assist tumors in developing powerful ‘generalist’ phenotypes as they evolve (Cook and Wrana, 2022). The mathematical model with which we study EMT population dynamics is phenomenological: capturing the rates of entry/exit between EMT states without transcriptional information or feedback signaling. It does not incorporate additional complexities such as reverse transitions or stochasticity. We have used external information from the biological properties of EMT to construct our mathematical model, and not obtained it purely from the cell dynamics observed in the data (Weinreb et al., 2018). Nonetheless, to compare relative transition rates, a simple three-compartment model seems reasonable to describe most conditions analyzed and fits both the inferred cell states (clusters) and the pseudotemporal dynamics during EMT. In the future, combining the cell population dynamics with a transcriptional EMT network (Hong, Watanabe, et al., 2015) to investigate the role of cell-cell communication (Rommelfanger et al., 2021) on the population dynamics of EMT could lead to additional insight – though additional data may be required for the transcriptional dynamics of such a model to avoid double dipping (Neufeld et al., 2024).

Canonical EMT states are defined by morphological features: epithelial cells adhere to each other with apical-basal polarity; mesenchymal cells are spindle-shaped, migratory, and lack cell-cell adhesion (Thiery et al., 2009). These morphological/adhesive properties cannot be fully captured by sequencing data alone. Moreover, multiple EMT gene lists (typically focusing on mesenchymal traits) have been proposed, with varying levels of agreement (Cook and Vanderhyden, 2022; Kinker et al., 2020; Puram et al., 2017; Tan et al., 2014; Yang et al., 2020). This variability in consensus genes also applies to epithelial genes, which can show tissue-specific heterogeneity. No single gene list can do justice to the heterogeneous paths of EMT, yet as we have shown, distinctive dynamic properties of EMT intermediate states can be captured by marker genes.

Genes predicted here as candidate markers of intermediate state EMT genes may serve as biomarkers of cells likely to metastasize and could be tested as predictors of clinical progression. In addition, such genes may mark for high-risk tumor cells prone to metastasis or recurrence, given the high metastatic potential of EMT intermediate state cells (Brown et al., 2022; Lüönd et al., 2021; Simeonov et al., 2021; Yu et al., 2013). More broadly, this study has shed new light on the plasticity of the EMT landscape and how it shapes the cell state transitions underlying cancer metastasis.

## AUTHOR STATEMENTS

### Author Contributions

**M.M.**: Conceptualization, software, methodology, investigation, formal analysis, writing— original draft, writing—reviewing & editing. **R.M.**: Investigation, writing—reviewing & editing. **E.T.R.T.**: Investigation, supervision, writing—reviewing & editing. **A.L.M.**: Conceptualization, software, methodology, investigation, funding acquisition, supervision, writing—original draft, writing—reviewing & editing.

### Funding Statement

This work was supported by the National Institutes of Health (R35GM143019) and the National Science Foundation (DMS2045327) (to A.L.M.).

### Conflict of Interest Statement

The authors declare no competing interests.

### Data Availability Statement

All code and data analysis associated with this study are released under an MIT license, available on GitHub: https://github.com/maclean-lab/dynamicEMT-genes. All raw sequencing data used in this study are publicly available on the Gene Expression Omnibus (GEO); see Methods for accession numbers.

## Notes

https://github.com/maclean-lab/dynamicEMT-genes

### Competing Interest Statement

The authors have declared no competing interest.

