## Supplementary Tables and Figures for "Modeling the dynamics of EMT reveals genes associated with pan-cancer intermediate states and plasticity"

---

#### Contents

**SUPPLEMENTARY TABLES** . . . . . . . . . . 1

**SUPPLEMENTARY FIGURES** . . . . . . . . . . 4

#### SUPPLEMENTARY TABLES

| Sample | Total counts threshold | Percent mitochondrial counts threshold | Percent ribosomal counts threshold | Leiden resolution |
|---|---|---|---|---|
| Mouse SCC, *in vivo* | 100,000 | 12% | – | 0.4 |
| HMLE stim. TGF-$\beta$ (8d) | 10,000 | 12% | 12% | 0.4 |
| HMLE stim. TGF-$\beta$ (8d) | 10,000 | 12% | 12% | 0.4 |
| HMLE stim. TGF-$\beta$ (10d) | 10,000 | 12% | 12% | 0.35 |
| HMLE stim. TGF-$\beta$ (10d) | 10,000 | 12% | 12% | 0.4 |
| HMLE stim. Zeb1 | 10,000 | 15% | – | 0.75 |
| HMLE stim. Zeb1 | 10,000 | 15% | – | 0.65 |
| A549 stim. TGF-$\beta$ | 40,000 | 8% | – | 0.4 |
| DU145 stim. TGF-$\beta$ | 50,000 | 8% | – | 0.4 |
| OVCA420 stim. EGF | 35,000 | 10% | – | 0.55 |
| OVCA420 stim. TGF-$\beta$ | 35,000 | 10% | – | 0.5 |
| OVCA420 stim. TNF | 40,000 | 10% | – | 0.45 |
| MCF10A stim. TGF-$\beta$ | 35,000 | 7% | – | 0.2 |

**Supplementary Table 1:** *Processing parameters for scRNA-seq data across each dataset.*

| Sample | $k1$,<br>Transition Rate $E \rightarrow I$ | $k2$,<br>Transition Rate $I \rightarrow M$ |
|---|---|---|
| HMLE stim. TGF-$\beta$ (8d) | 4.77 | 1.11 |
| HMLE stim. TGF-$\beta$ (8d) | 6.45 | 1.40 |
| HMLE stim. TGF-$\beta$ (10d) | 6.52 | 1.61 |
| HMLE stim. TGF-$\beta$ (10d) | 2.31 | 1.87 |
| HMLE stim. Zeb1 | 2.36 | 5.06 |
| HMLE stim. Zeb1 | 2.60 | 6.40 |
| A549 stim. TGF-$\beta$ | 2.38 | 3.16 |
| DU145 stim. TGF-$\beta$ | 1.50 | 1.57 |
| OVCA420 stim. EGF | 3.65 | 1.93 |
| OVCA420 stim. TGF-$\beta$ | 5.06 | 1.86 |
| OVCA420 stim. TNF | 2.07 | 1.68 |
| MCF10A stim. TGF-$\beta$ | 2.88 | 2.18 |

| Sample | $k1$,<br>Transition Rate<br>$E \rightarrow I1$ | $k2$,<br>Transition Rate<br>$I1 \rightarrow I2$ | $k3$,<br>Transition Rate<br>$I2 \rightarrow M$ |
|---|---|---|---|
| Mouse SCC, *in vivo* | 1.99 | 3.88 | 1.79 |

**Supplementary Table 2:** *Fitted mathematical $k_n$ model parameters for each scRNA-seq dataset. The in vivo mouse SCC dataset has two intermediate states with three $k_n$ parameters. All other datasets have one intermediate state with two $k_n$ parameters.*

| Gene | Int. DE | Int. DV | $k_1$ pos corr | $k_2$ neg corr |
|---|---|---|---|---|
| SFN | ● | ○ | ● | ● |
| NRG1 | ● | ● | ○ | ● |
| ITGB4 | ● | ● | ○ | ○ |
| ITGA6 | ● | ○ | ○ | ● |
| CBFB | ● | ○ | ● | ○ |
| FAM111A | ● | ○ | ○ | ● |
| LINC01503 | ● | ○ | ○ | ● |
| PLEK2 | ● | ○ | ○ | ● |
| IL4R | ● | ○ | ○ | ● |
| STK17A | ○ | ○ | ● | ● |
| CENPW | ○ | ○ | ● | ● |
| KRT18 | ○ | ○ | ● | ● |
| GJB3 | ○ | ○ | ● | ● |
| FHOD3 | ○ | ○ | ● | ● |

**Supplementary Table 3:** *Genes influencing intermediate EMT dynamics. DE denotes genes differentially expressed in intermediate states, and DV denotes genes with differential velocity in intermediate states. A positive $k_1$ correlation indicates faster $E \rightarrow I$ transition, while a negative $k_2$ correlation indicates slower $I \rightarrow M$ transition.*

### SUPPLEMENTARY FIGURES

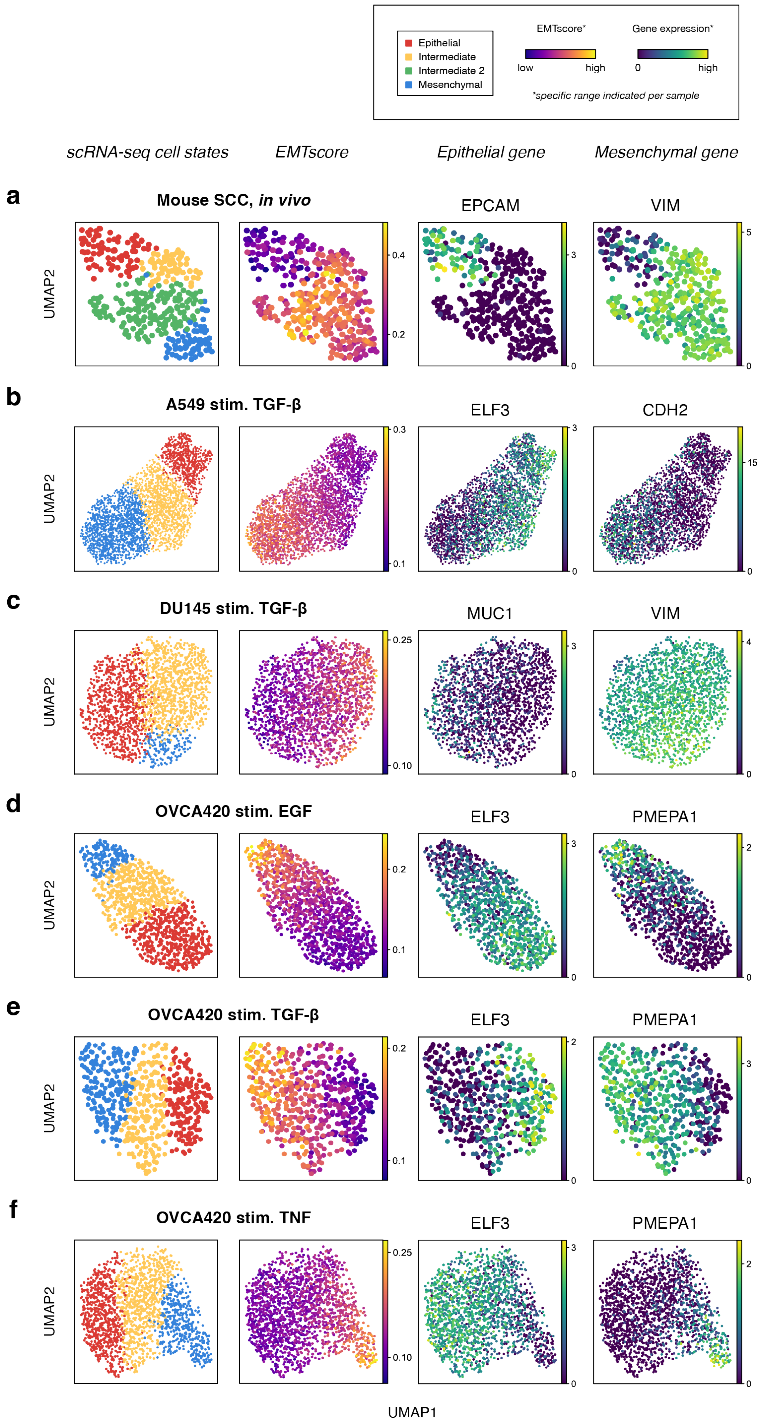


**Supplementary Figure 1:** ***a-f.*** *scRNA-seq data analysis of Pastushenko et al. (2018) and Cook and Vanderhyden (2020). Cell states were identified via Leiden clustering, EMT scores calculated with UCell, and representative epithelial and mesenchymal genes are shown.*

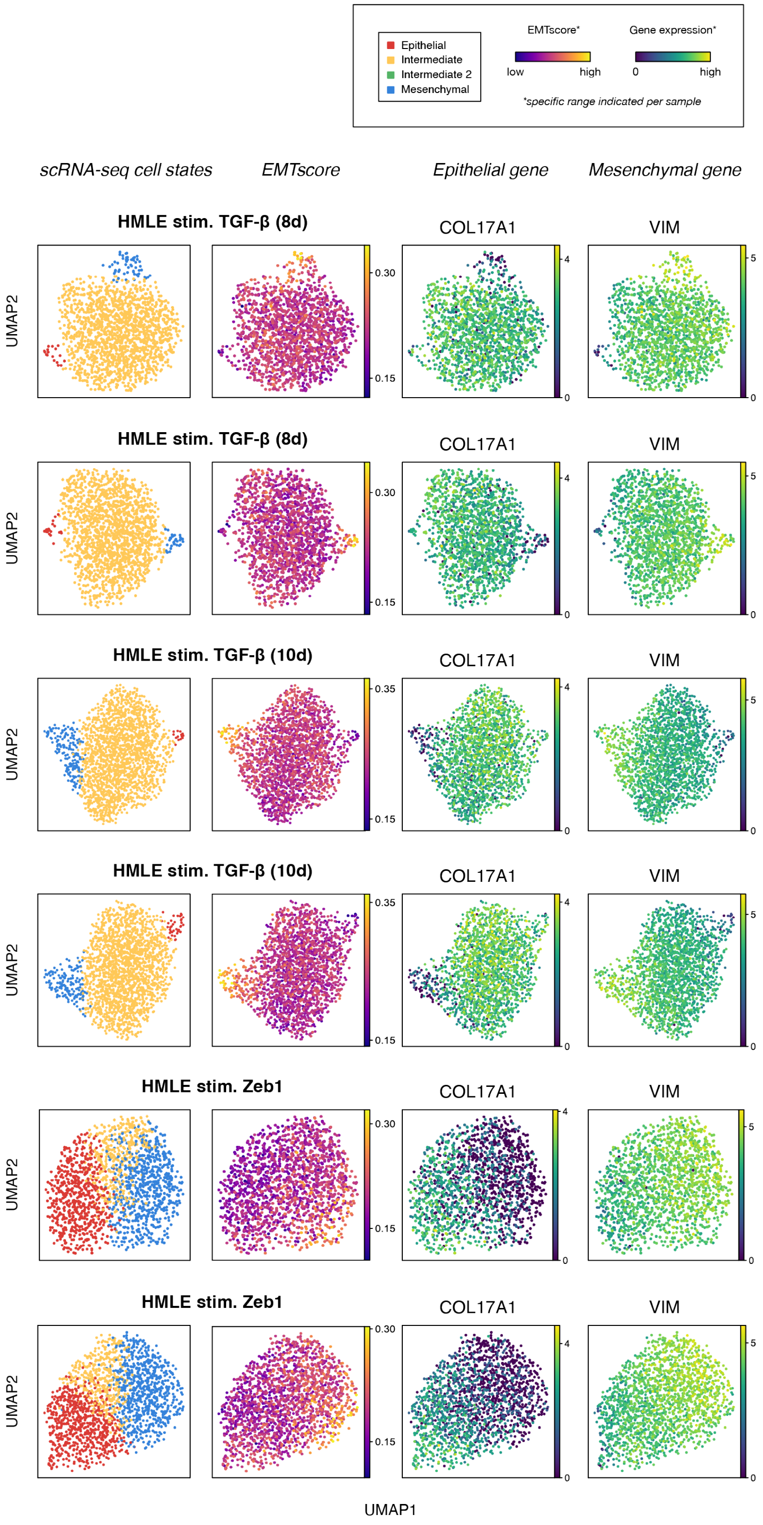


**Supplementary Figure 2:** ***a-f.*** *scRNA-seq data analysis of van Dijk et al. (2018). Cell states were identified via Leiden clustering, EMT scores calculated with UCell, and representative epithelial and mesenchymal genes are shown.*

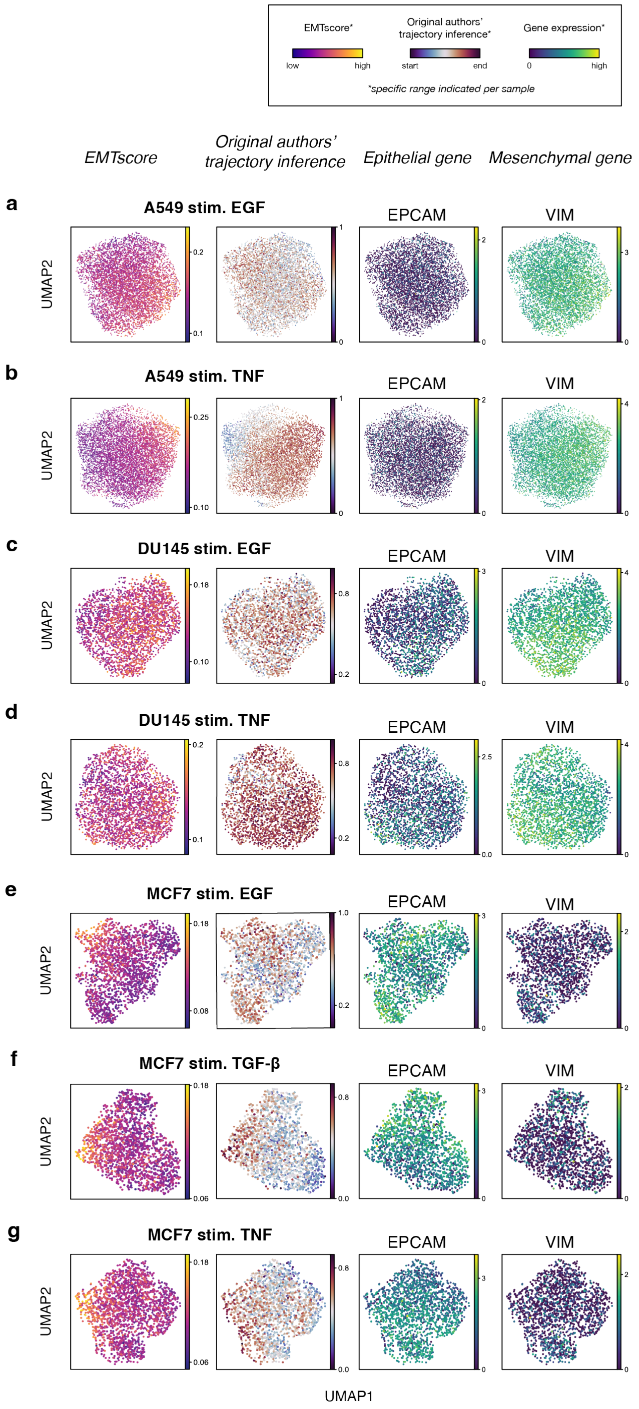


**Supplementary Figure 3:** ***a-g.*** *scRNA-seq data analysis of Cook and Vanderhyden (2020) for samples that did not exhibit a clear EMT and were excluded from the main analysis. EMT scores were calculated using UCell, and trajectory inference was obtained from the original publication. Representative epithelial and mesenchymal genes (EPCAM and VIM) are shown but did not align with distinct EMT cell states. The lack of EMT trajectory exemplified through both EMT scores and pseudotime values supports the lack of EMT in these samples.*

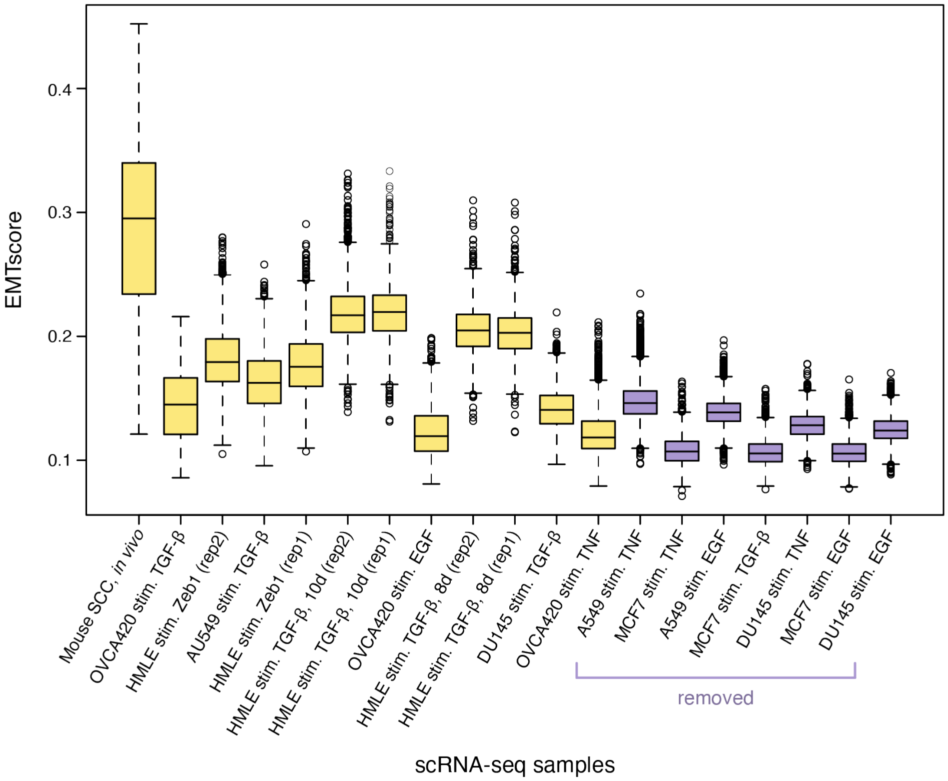


**Supplementary Figure 4:** *EMTscore distributions for all scRNA-seq datasets analyzed: mean and inter-quartile range of each dataset. The distributions are sorted by range from highest to lowest. Datasets were excluded if they did not exhibit EMT (see Supp. Fig. 3), not based on EMTscores. We see that the datasets that were removed consistently had the lowest EMTscore ranges, supporting the lack of EMT in these samples.*

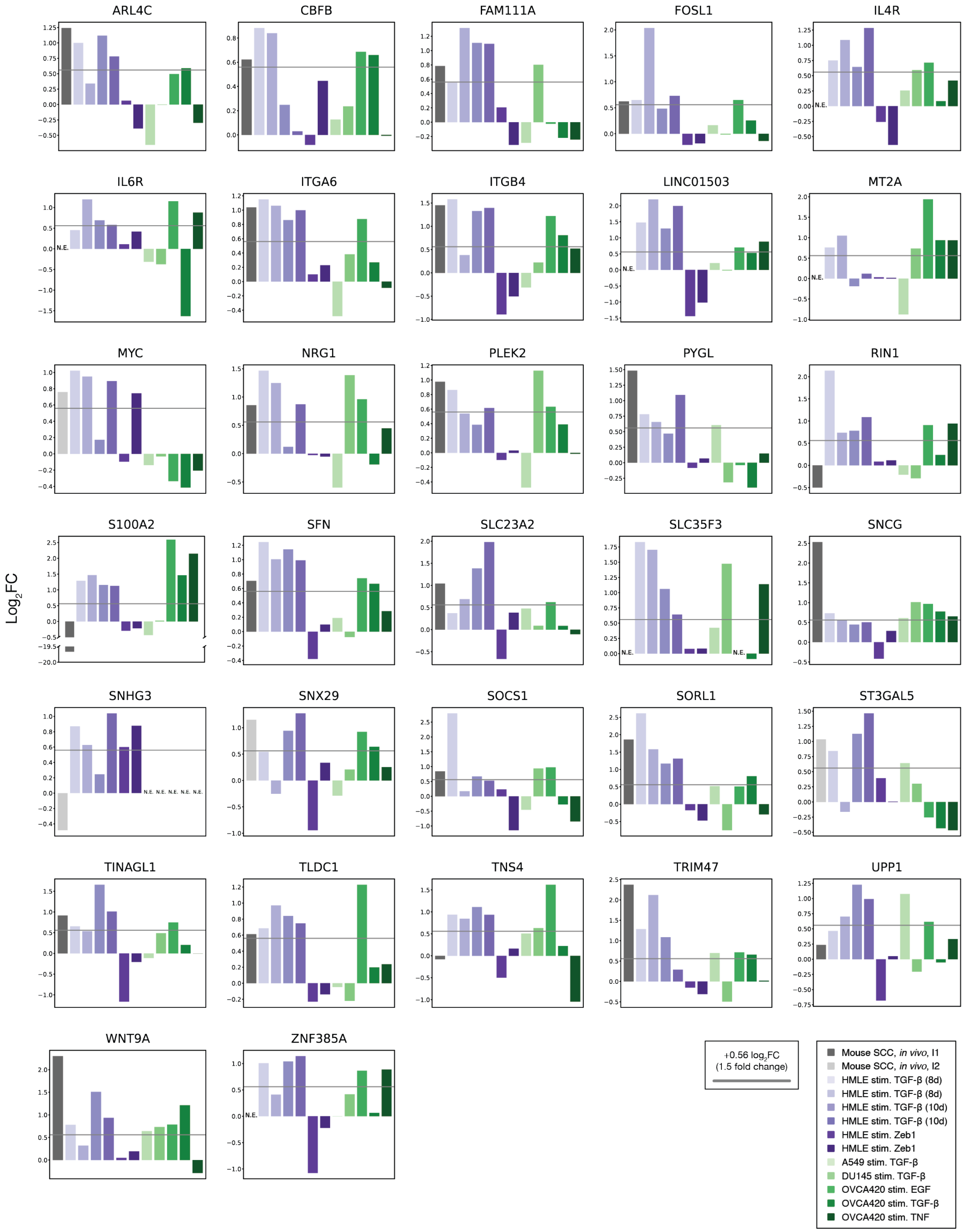


**Supplementary Figure 5:** *Genes associated with the intermediate EMT state, identified by $log_2FC$ upregulation in the intermediate state across scRNA-seq samples.*

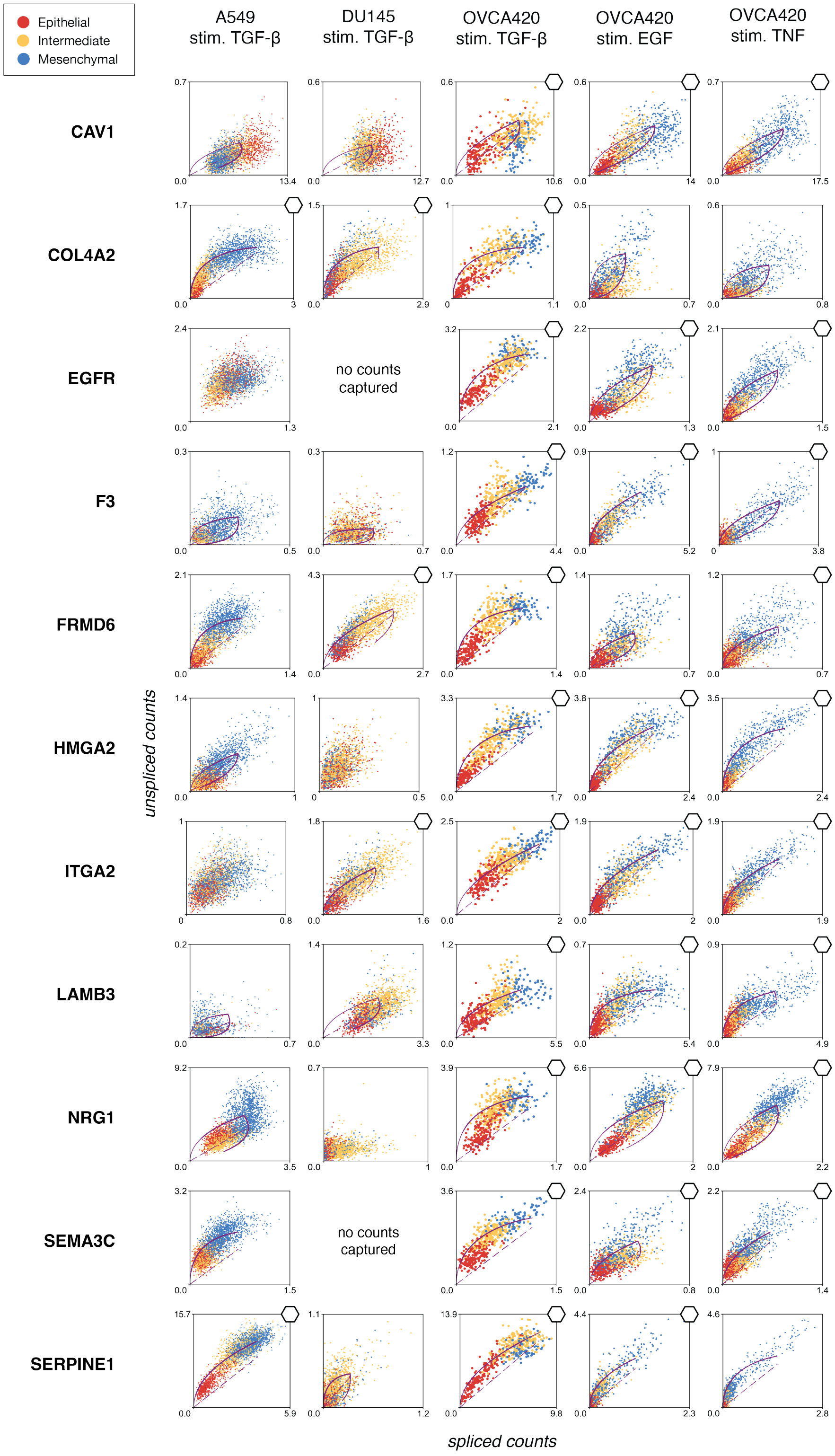


**Supplementary Figure 6:** *Genes with upregulated RNA velocity in the majority of intermediate EMT states across scRNA-seq samples from Cook and Vanderhyden (2020). Significant differential velocity for individual genes in specific samples indicated by ⬡.*

**a**

$$\frac{dE}{dt} = -k_1 E I_1$$

$$\frac{dI_1}{dt} = k_1 E I_1 - k_2 I_1 I_2$$

$$\dots$$

$$\frac{dI_n}{dt} = k_n I_{n-1} I_n - k_{n+1} I_n M$$

$$\frac{dM}{dt} = k_{n+1} I_n M$$

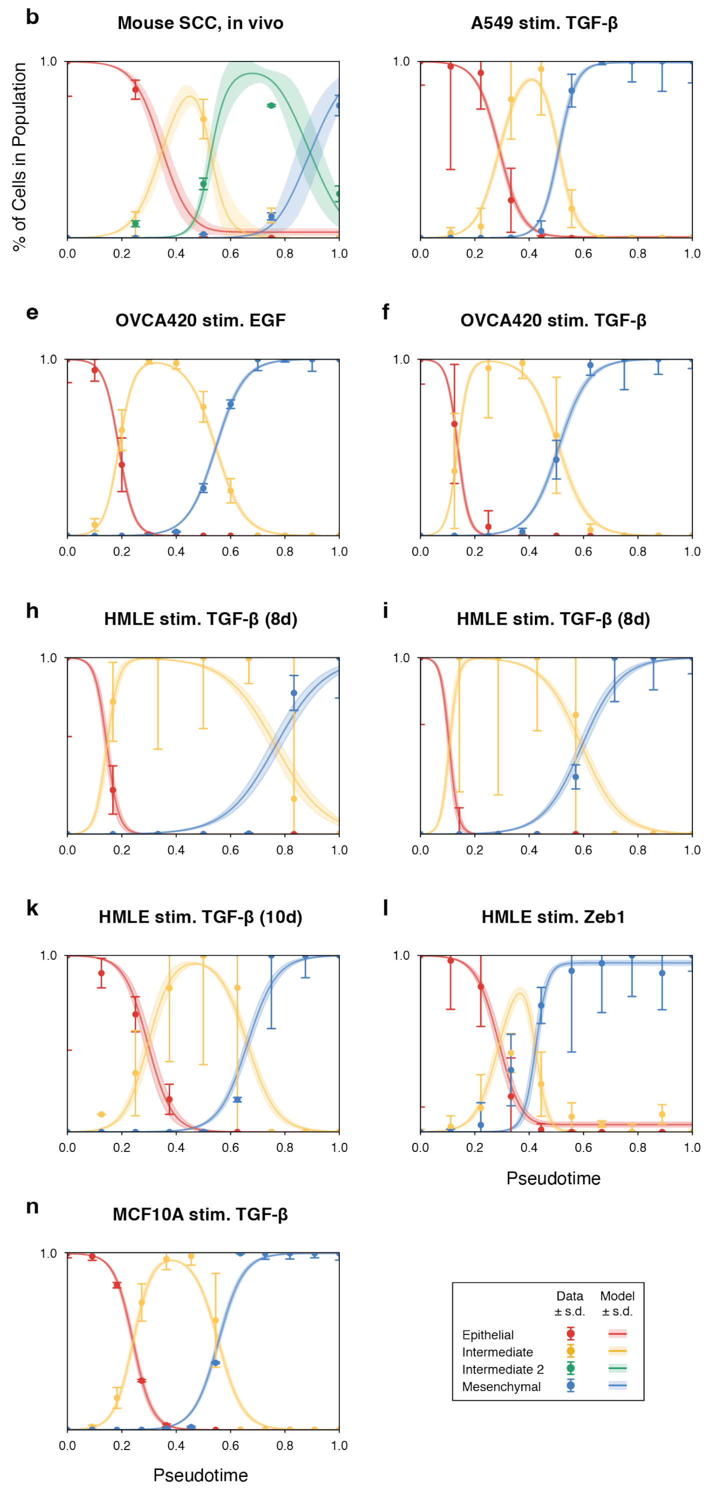


**Supplementary Figure 7:** ***a.*** *Generalized mathematical model for* $n$ *intermediate states.* ***b.*** *Model fits for each scRNA-seq sample following parameter inference, showing data vs. trajectory simulations with simulation parameters sampled from the posterior of each model.*

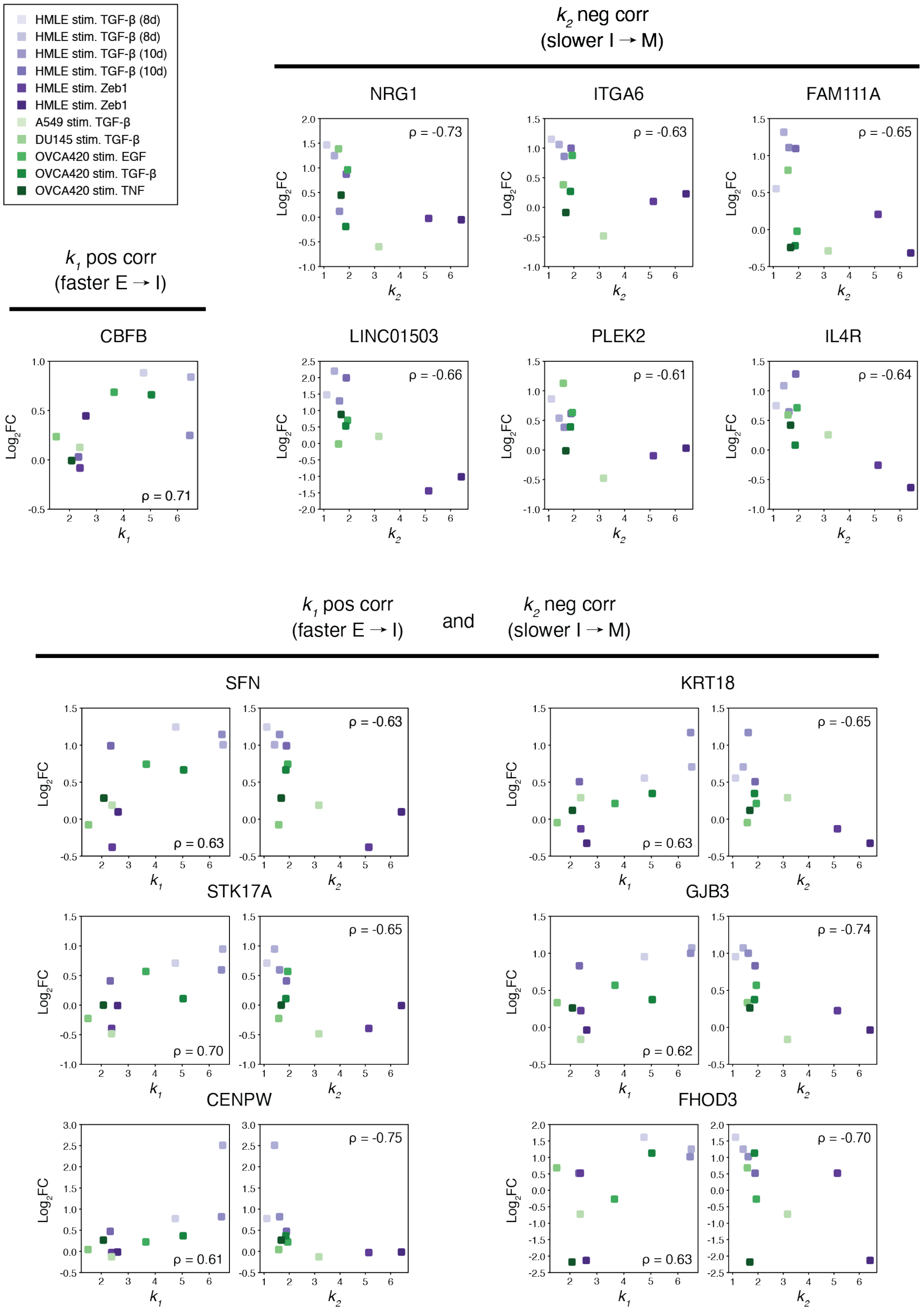


**Supplementary Figure 8:** *Intermediate gene $log_2FC$ correlated with fitted model parameters. Significant Spearman correlations ($p < 0.05$) were used to identify genes associated with stabilizing the intermediate state by increasing the transition rate into or decreasing the transition rate out of the intermediate state.*
